# Spectroscopic glimpses of the transition state of ATP hydrolysis trapped in a bacterial DnaB helicase

**DOI:** 10.1101/2021.04.08.438047

**Authors:** Alexander A. Malär, Nino Wili, Laura A. Völker, Maria I. Kozlova, Riccardo Cadalbert, Alexander Däpp, Marco E. Weber, Johannes Zehnder, Gunnar Jeschke, Hellmut Eckert, Anja Böckmann, Daniel Klose, Armen Y. Mulkidjanian, Beat H. Meier, Thomas Wiegand

## Abstract

The ATP hydrolysis transition state of motor proteins is a weakly populated protein state that can be stabilized and investigated by replacing ATP with chemical mimics. We present atomic-level structural and dynamic insights on a state created by ADP aluminum fluoride binding to the bacterial DnaB helicase from *Helicobacter pylori*. We determined the positioning of the metal ion cofactor within the active site using electron paramagnetic resonance, and identified the protein protons coordinating to the phosphate groups of ADP and DNA using proton-detected ^31^P,^1^H solid-state nuclear magnetic resonance spectroscopy at fast magic-angle spinning > 100 kHz, as well as temperature-dependent proton chemical-shift values to prove their engagements in hydrogen bonds. ^19^F and ^27^Al MAS NMR spectra reveal a highly mobile, fast-rotating aluminum fluoride unit pointing to the capture of a late ATP hydrolysis translation state in which the phosphoryl unit is already detached from the arginine and lysine fingers.

## Introduction

Adenosine triphosphate (ATP)-driven motor proteins play a key role in various cellular processes^1^. For example, motor proteins belong to the class of ATPases, which hydrolyze ATP into ADP (adenosine diphosphate) and inorganic phosphate to gain chemical energy allowing such enzymes to drive further chemical or mechanical events^2^. Structural insights into the functioning of these molecular machines are not straightforward to obtain, neither by X-ray crystallography, nor by cryo-electron microscopy or NMR spectroscopy due to the difficulty in trapping the intermediate catalytic states occurring during ATP hydrolysis. Diverse ATP analogues can be employed to mimic different stages of ATP hydrolysis as closely as possible^3,4^ which, in combination with molecular dynamics simulations^5^, can give mechanistic insights into complex biomolecular reaction coordinates. Of particular interest in unravelling the ATP hydrolysis reaction mechanism is the transition state of the phosphoryl (PO_3_^−^) transfer reaction (see Scheme 1 for a sketch of the limiting case of an associative ATP hydrolysis mechanism^6^). Metal fluorides have been found to mimic such states for structural studies, mostly using X-ray crystallography^7,8^. The number of deposited protein structures containing analogues such as AlF_4_^−^, AlF_3_ and MgF_3_^−^ has strongly increased in the past years^7^. AlF_4_^−^ forms together with the phosphate oxygen atom of ADP as well as an apical water molecule an octahedral complex mimicking the “in-line” anionic transition state of phosphoryl transfer, whereas AlF_3_ and MgF_3_^−^ form trigonal-bipyramidal complexes^7^. The formation of AlF_4_- or AlF_3_ is controlled by pH, the latter being favored at lower pH values^9^. However, some concern regarding their discrimination, *e.g.* distinction between AlF_3_ and MgF_3_^−^, has been raised and it could be shown by ^19^F NMR spectroscopy that some complexes, which were believed to contain AlF_3_, contain instead MgF_3_^− 9^. Similarly, in lower resolution X-ray structures (> 2.8 Å) AlF_3_ cannot be distinguished unambiguously from AlF_4_^− 9,10^.

**Scheme 1:**
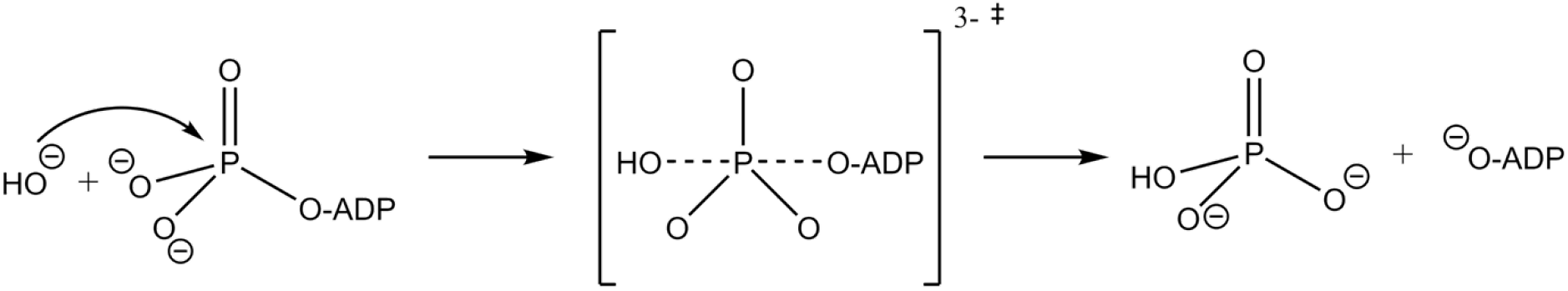
Sketch of the associative ATP hydrolysis mechanism with a trigonal-bipyramidal transition state. ‡ indicates the transition state.

We present magnetic-resonance approaches using EPR and solid-state NMR to obtain spectroscopic insights into the transition state of ATP hydrolysis which we trap for the oligomeric bacterial DnaB helicase from *Helicobacter pylori* (monomeric molecular weight 59 kDa) by using the transition-state analogue ADP:AlF_4_^−^. The motor domain of the helicase belongs to P-loop fold nucleoside-triphosphatases (NTPases), one of the largest protein families, which includes motor proteins like myosins, kinesins, and rotary ATPases. About 10-20% of genes in any genome encode for diverse P-loop fold NTPases^11^. In these enzymes, ATP or guanosine triphosphate (GTP) molecules are bound to the so-called Walker A motif GxxxxGK[S/T] of the signature P-loop of the motor nucleotide-binding domain (NBD)^12,13^. Bacterial DnaB helicases, which use the energy of ATP hydrolysis to unwind the DNA double helix, belong to the ASCE division of P-loop fold NTPases. The members of this division are characterized by an additional β-strand in the P-loop fold and a catalytic glutamate (E) residue next to the attacking water molecule^14–16^. Within the ASCE division, DnaB helicases are attributed to the RecA/F1 class^16^. Generally, P-loop fold NTPases need to be activated before each turnover because otherwise, they would promptly consume the entire cellular stock of ATP and GTP. As inferred from the comparative structure analysis of NTPases with transition state analogues, such as NDP:AlF_4_^−^ or NDP:MgF_3_^−^, the activation is mostly achieved by the insertion of a positively charged activating moiety (usually, an arginine or lysine “finger” or a potassium ion) between the α- and γ-phosphates^7,17–19^. As shown by MD simulations, linking of α- and γ-phosphates by the activating moiety leads to rotation of the γ-phosphate group yielding an almost eclipsed, hydrolysis-prone conformation of the triphosphate chain^20^. RecA-type ATPases, however, generally, make an exception; their activating moieties, which operate in a tandem, e.g. as Arg and Lys residues provided by the neighbouring subunit in a DnaB oligomer, can reach only the γ-phosphate group, but not the α-phosphate.

W-band electron-electron double resonance (ELDOR)-detected NMR (EDNMR)^21–24^ and electron-nuclear double resonance (ENDOR)^25,26^ allow for the positioning of the divalent metal ion within the active site by identifying nuclei in its vicinity. The native Mg^2+^ cofactor is replaced for such studies by the EPR-observable paramagnetic Mn^2+^ analogue^27^ (the biological functionality is maintained under such conditions^28^). ^19^F, ^27^Al and ^31^P nuclear resonances were observed among others in the EDNMR spectra, proving the binding-mode of ADP:AlF_4_^−^. The extracted ^31^P hyperfine coupling constants and the detection of ^19^F and ^27^Al nuclei in the proximity of the co-factor point to a coordination of the Mn^2+^ ion to the β-phosphate of ADP and AlF_4_^−^.

Solid-state NMR can identify the amino-acid residues involved in the coordination of the ATP analogue. Protons are of particular interest as their resonance frequencies can contain information regarding their engagement in hydrogen bonds. Fast magic angle spinning (MAS) nowadays provides sufficient spectral resolution for proton-detected side-chain studies^29^. Indeed, proton detection at fast MAS has become an important tool in structural biology in the past years for unraveling protein structures^30–39^, to characterize RNA molecules^40^ and protein-nucleic acid interactions^41–43^, and to address protein dynamics^44–50^. A key advantage of solid-state NMR is the straightforward sample preparation, which simply consists of sedimentation from solution into the solid-state NMR rotor without requiring crystallization steps^51–53^. We herein identify protein residues engaged in hydrogen bonding to the phosphate groups of nucleotides (ADP:AlF_4_^−^ and DNA) by (i) measuring high-frequency shifted proton resonances characteristic for hydrogen-bond formation^54^, (ii) probing spatial proximities in dipolar-coupling based proton-detected ^31^P,^1^H correlation experiments at fast MAS (105 kHz) and (iii) using the temperature-dependence of ^1^H chemical-shift values as a probe for hydrogen bonding, an approach well-known in solution-state NMR^55–57^, and recently extended to the solid state^58^. From a combination of (i)-(iii), key contacts between the ADP phosphate groups and residues located in the Walker A motif were identified, as well as two hydrogen bonds to the phosphate groups of the two DNA nucleotides. To complement our spectroscopic characterization of the ATP hydrolysis transition state, we performed ^19^F and ^27^Al MAS experiments to access information about bound AlF_4_^−^. The spectra indicate a fast rotation of the AlF_4_^−^ unit implying that the AlF_4_^−^ is not rigidified by coordinating protein residues indicating that the ADP:AlF_4_^−^ trapped state of DnaB possibly describes a late transition state, just after the bond fission, but before the release of the phosphate group from the catalytic pocket.

## Results

### EPR enables the positioning of the metal ion cofactor within the active site

Binding of ADP:AlF_4_^−^ to the protein is revealed in EDNMR experiments which employ the hyperfine couplings between a paramagnetic center and nearby nuclei to detect the latter. EDNMR has been used to characterize transition states of ATP hydrolysis, often in the context of ABC transporters for which such a state is successfully mimicked by ADP-vanadate^59,60^. Figure 1a shows the Mn^2+^ EDNMR spectrum of DnaB complexed with ADP:AlF_4_^−^ (red) compared to the reference spectrum of DnaB complexed only with ADP (cyan). While in both cases couplings to ^31^P nuclei are observed, additional peaks for ^19^F and ^27^Al are detected only for the ADP:AlF_4_^−^ bound state consistent with the presence of AlF_4_^−^ in the NBD of DnaB. To rule out that these correlations originate from the formation of the Mn^2+^:ADP:AlF_4_^−^ complex in solution, we recorded EDNMR spectra on a frozen control solution in the absence of protein and indeed we do not observe any ^19^F and ^27^Al resonances (purple spectrum in Figure 1a). Interestingly, in the presence of protein, two groups of ^31^P resonances are detected: a hyperfine-split doublet (denoted ^31^P^d^ in Figure 1a) and an unresolved doublet (denoted ^31^P^s^). Davies ^31^P Electron-Nuclear DOuble Resonance (Davies ENDOR)^25^ experiments were performed on the Mn^2+^-containing protein complex (Figure 1b) to extract the hyperfine tensor *A* of the doublet. Line-shape simulations yield a large *A*_iso_ value of 4.7 MHz (for all ^31^P hyperfine tensor parameters extracted from the spectrum see Table S1). This value is in agreement with published values for an ADP:Mn^2+^ complex in which the Mn^2+^ ion binds symmetrically to the two ADP phosphate groups^61^. Mims ENDOR^26^ experiments were performed to detect the small *A*_iso_ value of the in EDNMR unresolved doublet which is determined to be 0.3 MHz (see Figure 1c). Mims ENDOR measurements on the control solution did not show this doublet. We assign the large *A*_iso_ value (4.7 MHz) to ^55^Mn-^31^Pβ and the small *A*_iso_ value (0.3 MHz) to ^55^Mn-^31^Pα hyperfine couplings indicating that the Mn^2+^ ion is located much closer in space to the Pβ atom of ADP than to the Pα atom. This assignment is supported by Density Functional Theory (DFT) calculations of the hyperfine coupling tensors performed on small clusters mimicking the Mn^2+^ coordination sphere extracted from the available crystal structures of SF4 helicases (the same super-family to which *Hp*DnaB belongs, PDB accession codes 4ESV: *Bst*DnaB:ADP:AlF_4_^−^:DNA^62^ and 4NMN: *Aa*DnaB:ADP^63^, *vide infra*) although it has to be noted that the uncertainty in the exact metal ion position due to insufficient resolution of the electron density and the initial presence of Ca^2+^ instead of Mg^2+^ in the 4ESV structure might be significant and influence the results of the calculations (Table S1 and Figure S1 in the Supplementary Materials Section).

**Figure 1:**
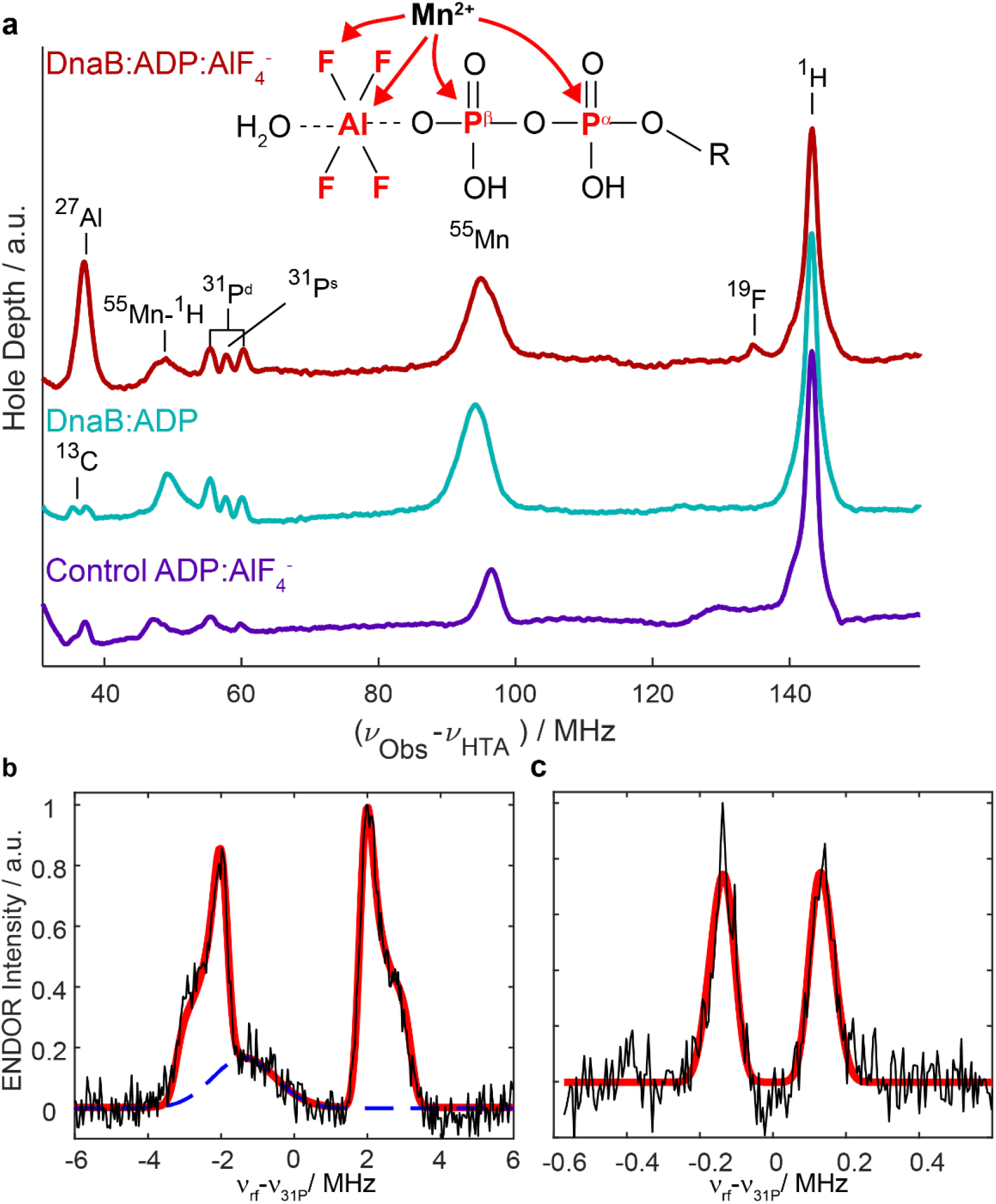
EPR characterizes binding of the metal ion cofactor and ADP:AlF_4_^−^ to DnaB. **a** EDNMR highlighting hyperfine couplings and thus proximities between the Mn^2+^ metal center and surrounding nuclei measured for DnaB:ADP:AlF_4_^−^ (red) and DnaB:ADP (cyan), as well as for a control solution containing only Mn^2+^:ADP:AlF_4_^−^ in the same buffer used for the protein sample (purple). The assignments of the peaks to the nuclear resonance frequencies are shown. Parts of the cyan spectrum are reproduced from reference^60^. ^31^P Davies ENDOR (**b**) and Mims ENDOR (**c**) recorded on DnaB:ADP:AlF_4_^−^. The red lines represent line shape simulations using EasySpin^64^ based on *A*_iso_-values of 0.3 and 4.7 MHz (for all parameters see Table S1). The broad background peak in (**b**) is most likely a third harmonic of one of the Mn^2+^ hyperfine lines and was removed for the fitting.

### Hydrogen bonds to the phosphate groups of ADP and DNA nucleotides identified by fast MAS experiments

Solid-state NMR experiments on DnaB complexed with ADP:AlF_4_^−^ and single-stranded DNA (a polythymidine stretch with 20 DNA nucleotides was used^65^) allow a direct view into the NBD. Figure 2a shows the previously reported ^31^P-detected cross-polarization (CP)-MAS spectrum of DnaB in complex with ADP:AlF_4_^−^ and DNA (see Figure 2b for the atomic numbering) recorded at 17 kHz MAS^66^. Two narrow resonances are detected for both, the Pα and Pβ of ADP (at −6.0 and −7.1 ppm, respectively) as well as for the DNA phosphate groups (at 0.5 and −1.1 ppm). The latter observation reflects that two DNA nucleotides bind to one DnaB monomer which is characteristic for SF4-type helicases^42,66^. Proton-detected NMR experiments at fast MAS frequencies (> 100 kHz) allow the identification of protons engaged in hydrogen bonds requiring only small amounts of protein in the order of 0.5 mg. The ^1^H NMR chemical-shift value serves as a sensitive indicator for the formation of hydrogen bonds: a de-shielding effect is observed if protons are engaged in such interactions^42,43,54,67^. However, the chemical shift alone is not a sufficient criterion to prove hydrogen bonding. We therefore extend the experimental approaches to directly detecting such interactions by the presence of through-space ^31^P,^1^H dipolar couplings in hPH correlation experiments at 105 kHz MAS. The hPH spectra were recorded with two different ^1^H-^31^P CP contact times (1.5 and 3.5 ms) on a ^13^C,^15^N uniformly labeled, deuterated and 100% back-exchanged sample of DnaB in which the ADP and the DNA remained at natural abundance. Note that this deuterated version of the protein has been chosen over a fully-protonated sample to increase the intrinsically rather low signal-to-noise ratio in such a large protein due to the narrowing of the proton resonances by roughly a factor of three attributed to the dilution of the proton dipolar network (see Figure S2 for the proton line-widths determined for a deuterated and fully protonated sample)^68^. Figure 2c shows the rather sparse 2D hPH correlation spectrum (with 3 ms CP contact time) of the DnaB complex and indeed protein-phosphate correlations to all four ^31^P resonances observed in Figure 2a are visible. The CP-based hPH experiment proves spatial proximities between proton nuclei in the vicinity of the phosphate groups. An INEPT-based experiment transferring polarization directly over the hydrogen bond via the *J*-couplings (typical ^2^*J*(^31^P-^1^H) values are in the order of 3 Hz^69,70^) was not successful due to a too short proton transverse relaxation time compared to the required INEPT transfer delay period (see Figure S3). The resonance assignments shown in Figure 2c and Figure S4 (CP contact time of 1.5 ms) were obtained using the deposited proton chemical-shift values (BMRB accession code 27879). The hPH spectra reveal intense signals and thus spatial correlations between Pβ of ADP and S206, G208, K209 and T210, all located in the conserved Walker A motif of the P-loop in the motor domain of the helicase^71^. Note that for all mentioned amino acids correlations to the backbone amide protons are observed, except for K209 for which additional sidechain Hζ protons are detected. For the Pα resonance of ADP only weak correlations are observed, the strongest one to S211 and an unassigned resonance, possibly an arginine residue which has been detected in previous NHHP experiments^42^ (R242 or the “arginine finger” R446 from a neighboring DnaB subunit). The main difference in the spectrum recorded at shorter CP contact times (Figure S4) is that correlations to the ADP and DNA protons (sugar and base) present in the spectrum recorded at 3.5 ms contact time (highlighted in light red and green in Figure 2c) are absent. It is important to note that the herein described hPH experiments appear to be much more selective for detecting direct coordination partners than the previously described ^1^H-^1^H spin-diffusion based NHHP and CHHP experiments^72^ and possibly also TEDOR experiments^73^, thereby providing a more detailed picture of the local geometry around the phosphate groups of ADP and DNA^4,74^ than reported previously^42^. The hPH spectrum in Figure 2c also contains important information regarding the DNA coordination. Actually, only two intense backbone amide correlations to the two DNA phosphate groups, D374 in case of P1 and G376 in case of P2 are observed. Together with our previous observation of K373 forming a salt-bridge to P2 via the lysine sidechain, only three contacts seem to coordinate the DNA in this molecular recognition process.

**Figure 2:**
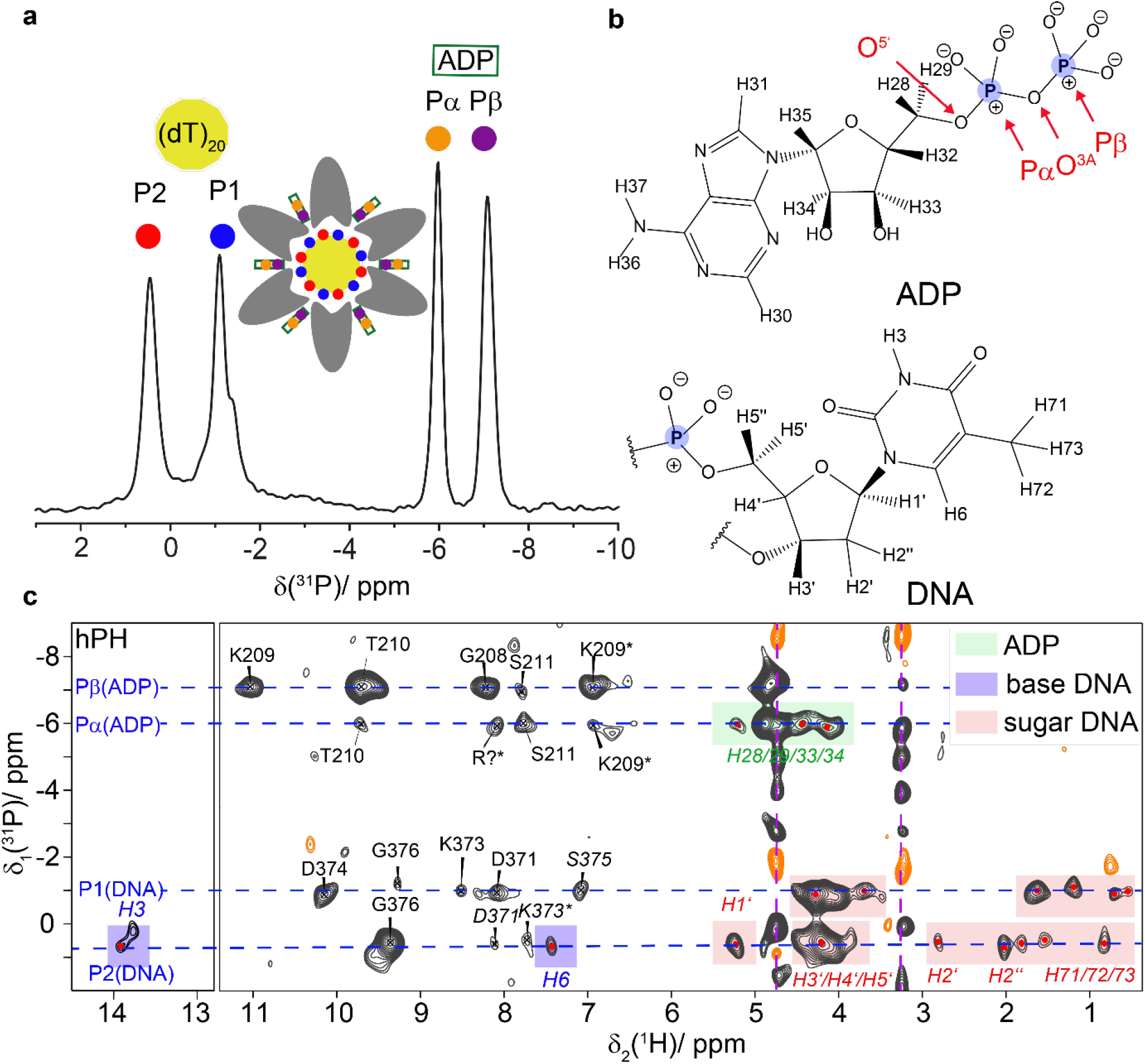
ADP and DNA recognition in DnaB highlighted by phosphorus-proton contacts identified at fast MAS. **a** ^1^H→^31^P (hP) CP-MAS spectrum of DnaB:ADP:AlF_4_^−^:DNA adapted from reference^66^ showing the resonance assignments of the DNA and ADP phosphate groups. The shoulder in the ^31^P resonance at ~−1.4 ppm possibly results from rigidified DNA nucleotides, which are, however, not coordinating to DnaB. **b** Chemical structures of ADP and DNA (thymidine) molecules including the numbering of proton atoms following the convention of the BMRB database (DNA) and the recent IUPAC recommendations for nucleoside phosphates^75^. Phosphorus atoms are highlighted in blue. **c** CP-based hPH correlation spectrum (CP contact time 3 ms) recorded on DnaB in complex with ADP:AlF_4_^−^ and DNA at 20.0 T external magnetic field and 105 kHz MAS. The protein resonance assignment is taken from reference^42^ (BMRB accession code 27879). Regular-printed residue labels: Chemical-shift deviation to reported proton shifts < 0.05 ppm. Italic-printed residue labels: Chemical-shift deviation to assigned proton shifts ≥ 0.05 ppm. All proton shifts are assigned to amide backbones, except the ones indicated by a * which are associated to side-chain atoms. Correlations between the phosphate groups and ADP or DNA are highlighted in green and light red/purple, respectively. The assignments of the DNA proton resonances are based on average chemical-shift values reported in the BMRB data base (www.bmrb.wisc.edu/). The pink dashed lines highlight signals from insufficiently suppressed DNA in solution.

The high-frequency shifts of their amide protons and their spatial proximity to the phosphate ADP group already point to the engagement of K209 and T210 in hydrogen bonding as discussed above. To further verify this, we determined the temperature dependence of their chemical shifts between 294 K and 302 K (sample temperatures, see Materials and Methods Section). Due to their characteristic chemical shifts (and thus their isolated position in the 2D fingerprint spectrum) the temperature-dependences could be directly extracted from 2D CP hNH experiments. It is well-known from solution-state NMR that the chemical shifts of protons in strong intramolecular hydrogen bonds experience only a weak temperature dependence^56,57^ as recently also shown by solid-state NMR^58^. However, for protons in rather weak hydrogen bonds, the resonances become significantly more shielded upon increasing the temperature, due to an increase in the average hydrogen-bond length. Figure 3 shows the temperature dependence for residues identified in the hPH spectra (left column). Indeed, K209 and T210, previously identified as forming hydrogen bonds to the Pβ of ADP, show an almost vanishing temperature coefficient (slope of the corresponding linear regression). Similar values are found for D374 and G376 (Figure S5) assumed to be involved in DNA coordination. In contrast, Figure 3 (right column) shows resonances associated with a larger temperature coefficient thus not being involved in hydrogen bonds (for all extracted temperature coefficients see Figure S5).

**Figure 3:**
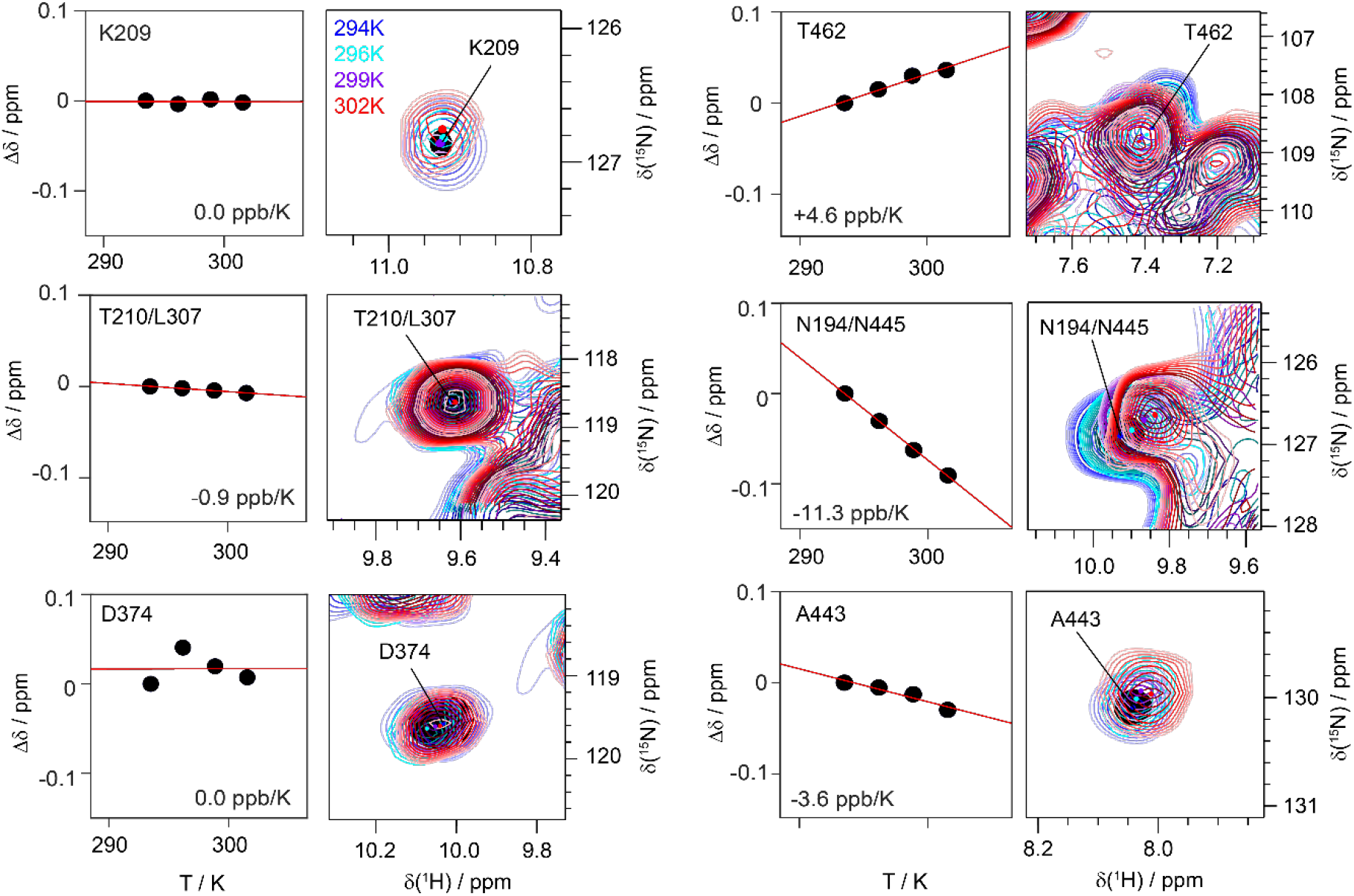
Temperature-dependent proton chemical-shift values as indicators for hydrogen bond formation. Residue-specific temperature coefficients and corresponding temperature-dependent hNH spectra (based on two ^1^H,^15^N CP steps) recorded at 20.0 T with a spinning frequency of 100 kHz for deuterated and 100 % back-exchanged DnaB complexed with ADP:AlF_4_^−^ and DNA. Temperature-dependent chemical-shift deviations (black circles) are referenced to the corresponding value at 294 K sample temperature.

### Solid-state NMR shows that the AlF_4_^−^ unit is highly mobile

The AlF_4_^−^ unit can be detected in ^19^F- and ^27^Al-detected MAS experiments. Figure 4a displays the ^19^F MAS spectrum of DnaB:ADP:AlF_4_^−^ in the presence and absence of DNA. Interestingly, only one ^19^F resonance line at around −146 ppm is detected for the AlF_4_^−^ group pointing to a fast chemical-exchange process (for the ^19^F spectrum in the absence of protein see Figure S6). The resonance assignments displayed in Figure 4a are based on reported solution-state NMR assignments^76,77^. A similar chemical-exchange process has also been observed for the RhoA/GAP:GDP:AlF_4_^−^ complex^78^, for the GTPase hGBP1^79^ and for the motor protein myosin^76^ in solution-state ^19^F NMR experiments.

**Figure 4:**
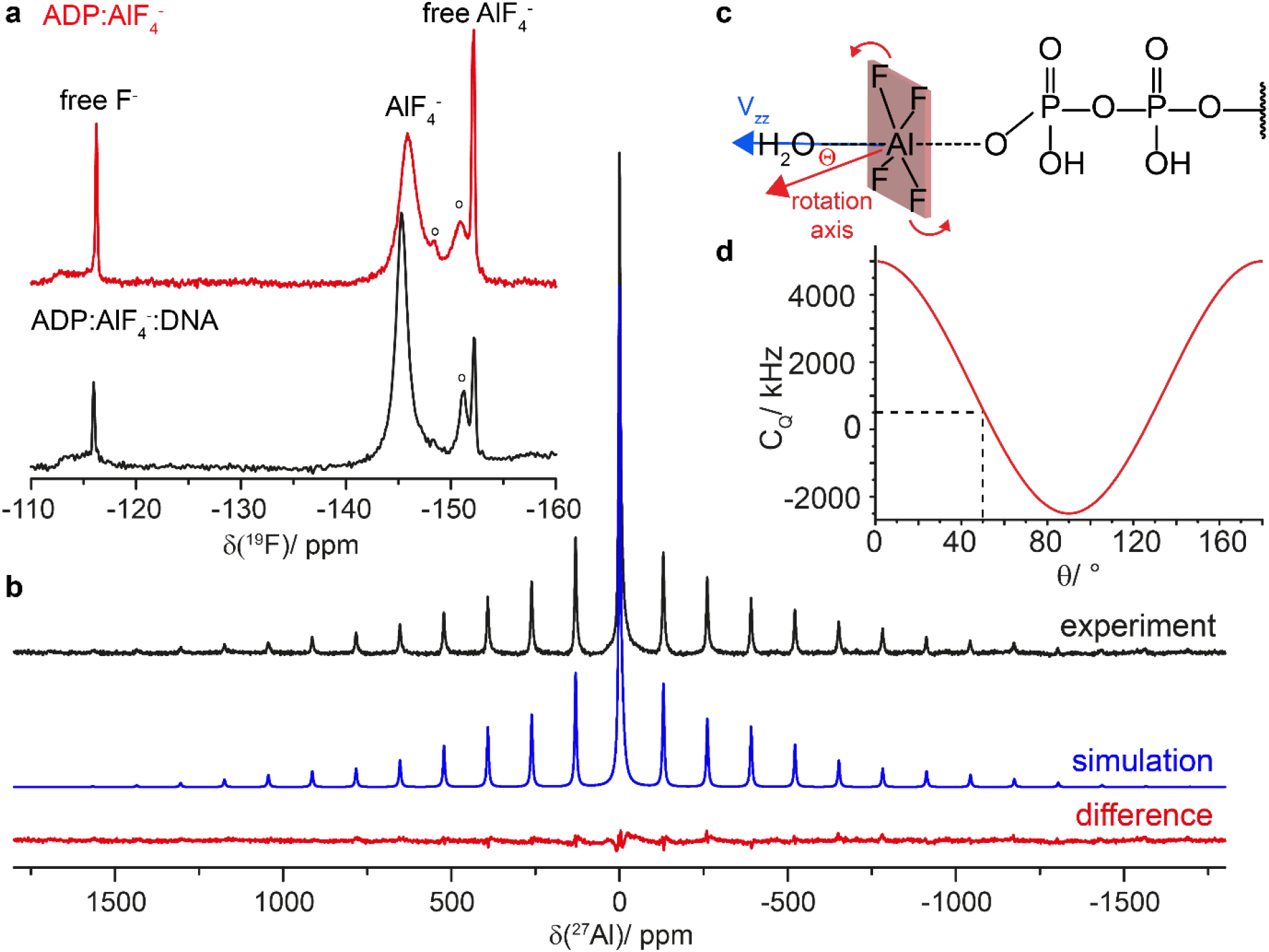
The AlF_4_^−^ species bound to DnaB is rotating. **a**^19^F MAS-NMR spectra recorded at 14.0 T with a MAS frequency of 17.0 kHz and with the EASY background suppression scheme^83^. Spectra were acquired on DnaB:ADP:AlF_4_^−^ in the presence and absence of DNA; “o” indicate precipitated AlF_x_(OH)_6-x_ species. **b** ^27^Al MAS-NMR spectrum of DnaB:ADP:AlF_4_^−^:DNA recorded at 11.74 T with a spinning frequency of 17.0 kHz (black) and corresponding line shape simulation using DMFIT^84^ assuming C_Q_(^27^Al)= 570 kHz, η_Q_(^27^Al)= 0.98, Δ_σ_(^27^Al)=−186 ppm and η_σ_(^27^Al)= 0.64. The central resonance is fitted with two additional Lorentzian lines possibly originating from aluminum hydroxyl fluorides in solution^77^. The difference spectrum is shown in red. **c** Schematic illustration of the rotation of the AlF_4_^−^ molecule. θ describes the angle between the rotation axis and the principal component (*V*_zz_) of the ^27^Al electric field gradient pointing along the O^…^Al^…^O axis. **d** Calculation of the effective ^27^Al quadrupolar coupling constant according to 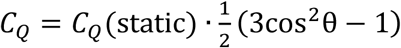, assuming a static C_Q_ value of 5 MHz (AlF_4_O_2_ species in AlF_x_(OH)_3-x_ ·H_2_O as taken from reference ^81^). Fast rotation of the AlF_4_^−^ unit on the NMR time scale is assumed in this calculation.

The ^27^Al satellite transition NMR spectrum (SATRAS, Figure 4b) of the sideband family is observed at δ_iso_= −0.2 ppm pointing to an octahedral coordination geometry of the ^27^Al nucleus^80^. The spectrum allows to extract the quadrupolar coupling constant (C_Q_) which amounts to only ~570 kHz. The central m = ½ ⟷ m = −½ transition is observed at a similar resonance shift indicating a small contribution of the second-order quadrupolar shift. The C_Q_-value is significantly lower than expected for a six-fold oxygen/fluorine coordinated aluminum species. C_Q_-values for crystalline aluminum hydroxyfluorides are typically in the order of 5 MHz^81^. We attribute this effect to a rotation on the NMR time scale of the AlF_4_^−^ unit around an axis inclined by an angle θ with respect to the direction of the principal component of the electric field gradient tensor (*V*_zz_, see Figure 4c). The angle θ must be close, about 5-10°, to the magic angle (54.7°) leading to the significant reduction of the anisotropy of the quadrupolar interaction (see Figure 4d). Similar observations were made for the DnaB complex in the absence of DNA (see Figure S7). Alternatively, the reduction of the quadrupolar interaction could be achieved by a rotational diffusion process, in which the angle θ varies randomly and is on average close to the magic angle. Note that the coordination of AlF_4_^−^ to the β-phosphate of ADP is also reflected in a low-frequency shift of the corresponding ^31^P ADP resonance (−4.5 ppm compared to the ADP-bound state^66^) which is a similar trend than observed for aluminium phosphate gels and glasses^82^.

The rotational motion or even diffusion of this unit (with a correlation faster than the inverse quadrupolar coupling constant) reflects the absence of tight binding either to the protein (e.g. via hydrogen bonds to the fluorine atoms) or to the metal ion cofactor. The ^27^Al isotropic chemical-shift value of close to 0 ppm is characteristic for an octahedrally coordinated Al-species, in our case most likely formed by four fluoride ligands, one oxygen ligand from the ADP phosphate backbone and one water molecule originating from an “in-line” geometry of phosphoryl transfer^7^ or the catalytic glutamate as observed in the *Bst*DnaB structure^62^.

### Homology modelling points to a free rotating AlF_4_^−^ detached from lysine and arginine fingers in SF4 helicases

We performed homology modelling based on the available bacterial helicase structures to investigate whether the dynamic behaviour of the AlF_4_^−^ moiety could be related to the activation mechanism of RecA NTPases. Although the crystal structure of the *Hp*DnaB dodecamer is available (PDB accession code 4ZC0^85^), its low resolution of 6.7 Å and the absence of either DNA or of bound nucleotides prevents its use for modelling the ADP:AlF_4_^−^ interactions in the catalytic site of a DNA-bound protein. Therefore, we reconstructed the mechanism of the activation from analysis of the DNA- and Ca^2+^:GDP:AlF_4_^−^-containing *Bst*DnaB structure (PDB accession code 4ESV, resolution 3.2 Å)^62^. In the *Bst*DnaB structure, the Ca^2+^:GDP:AlF_4_^−^ moieties are bound to five out of six catalytic centres (Figure S8). Furthermore, the positions and orientations of the AlF_4_^−^ moieties differ among the five catalytic sites (Figure S8). By considering these different configurations as mimics of different reaction intermediates, the reaction steps could be reconstructed in the following way. An apically positioned water molecule “attacks” the γ-phosphate group in the transition state^17^. In numerous P-loop fold NTPases this step manifests itself in formation of transition-state analogue complexes NDP:AlF_4_^−^:H_2_O_cat_ or NDP:MgF_3_^−^:H_2_O_cat_^7,8^. However, such a state with an apically placed H_2_O_cat_ is not observed in any of the five AlF_4_^−^-containing sites of the *Bst*DnaB structure. Therefore, in Figure 5a, we model this transition state using two structures as templates, the ADP:AlF_4_^−^:H_2_O_cat_ structure from the ABC-NTPase of the *E. coli* maltose transporter (which belongs to the same ASCE division as DnaB), as well as the whole structure of the closely related ADP:AlF_4_^−^-containing RecA of *E. coli* (with the anticipated catalytic water molecule unresolved). As seen in Figure 5a, the activating Arg and Lys residues in RecA form H-bonds with two fluorine atoms of AlF_4_^−^ (blue dashed lines). Comparison of the AlF_4_^−^ positions in different monomers of the *Bst*DnaB structure, as shown in Figures 5b, 5c and S8, suggests that Arg and Lys residues are able, together, to rotate and pull the γ-phosphate group, which is mimicked by AlF_4_^−^ in Figures 5b, c. While in Figure 5a and 5b the activating Lys and Arg residues are H-bonded to AlF_4_^−^, its further movement away from the nucleotide, as seen in Figure 5c, leads to the weakening of H-bonds or even their entire dissociation (note the longer distances indicated in Figure 5c), possibly yielding an almost unbound AlF_4_^−^ unit tilted relative to its catalytic position (compare Figure 5c with Figure 5a) in agreement with our solid-state NMR observations of a nearly freely rotating AlF_4_^−^ moiety.

**Figure 5:**
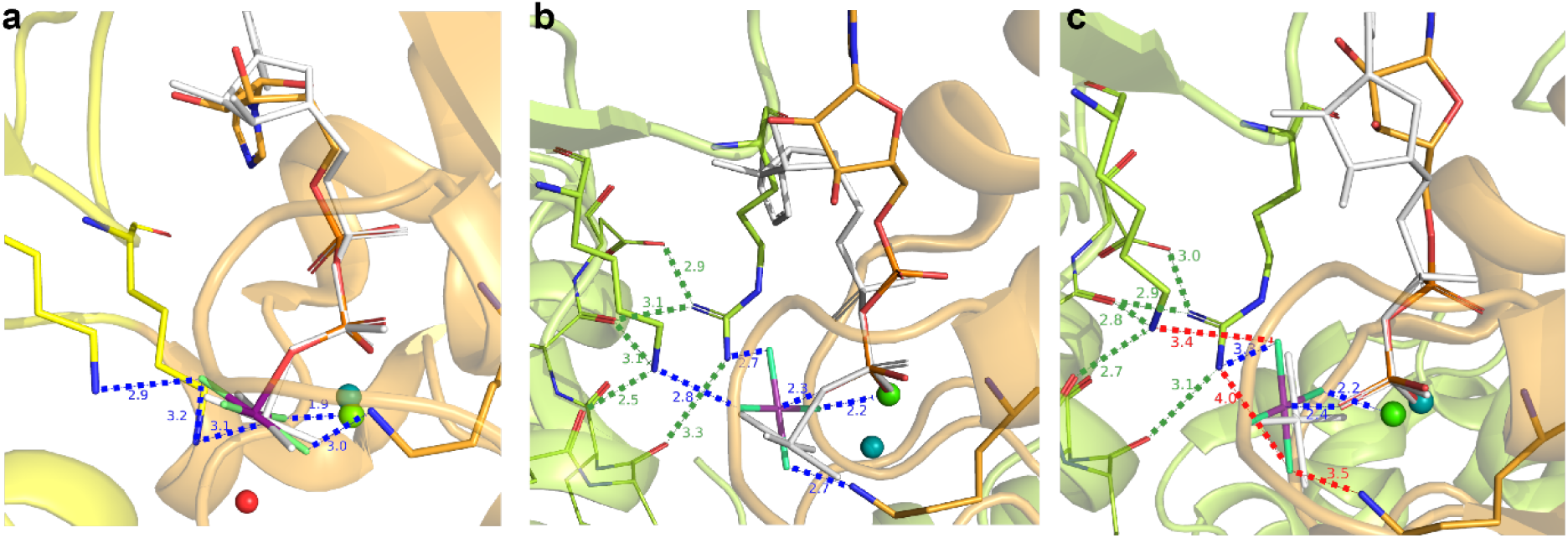
The transition-state analogue AlF_4_^−^ can adopt different orientations in diverse P-loop fold ATPases of the ASCE division. **a** AlF_4_^−^ binding in RecA from *E. coli* (PDB accession code 3CMW^86^). The P-loop domain (NBD domain) is shown in orange, the domain that provides the activating “fingers” is colored yellow. The magnesium ion is shown as a green sphere. To show the catalytic water molecule H_2_O_cat_, the ADP:AlF_4_ ^−^ complex is superimposed with the structure of the ADP:AlF_4_^−^:H_2_O_cat_ complex from the ABC ATPase of the maltose transporter MalK (see PDB accession code 3PUW^87^. The ADP molecule and AlF_4_^−^ of MalK are shown in white, H_2_O_cat_ as a red sphere, Mg^2+^ as a teal sphere. NDPs were superimposed using atoms O^3A^, P^B^ and O^3B^ (see Figure 2b for the atom notation used for ADP) in Pymol^88^. **b** Coordination of AlF_4_^−^ in the *Bst*DnaB structure (see Figure S8, PDB ID 4ESV and reference^62^). The nucleotide-binding chain C is colored orange, the activating chain B is colored light green. To show the displacement of AlF_4_^−^, the GDP:AlF_4_^−^ complex is superimposed, as described for panel **a**, with ADP:AlF_4_^−^ bound to the RecA protein (PDB ID 3CMW, see panel **a**). The H-bonds formed by AlF_4_^−^ are shown in blue, the additional interactions that stabilize the position of activating sidechains of K418 and R420 are shown in green. The AlF_4_^−^ moiety is twisted in comparison to the transition-state-mimicking complex shown on panel **a**. **c** Coordination of AlF_4_^−^ in the structure of *Bst*DnaB (see Figure S8, PDB ID 4ESV^62^). The nucleotide-binding chain F is colored orange, the activating chain E is colored light green. To show the further displacement of AlF_4_^−^, the GDP:AlF_4_^−^ complex of subunit F is superimposed, as described for panel **a**, with the same complex bound to subunit C, which is white colored (see panel **b**). Bonding interactions that are observed for the GDP:AlF_4_^−^ complex trapped at the B/C interface (see panel **b**), but not in this complex trapped at the E/F interface are shown as red dashed lines.

## Discussion

EPR experiments allow the localization of the metal ion co-factor within the NBD. In the transition state of ATP hydrolysis for *Hp*DnaB, the Mn^2+^ ion is coordinating to the β-phosphate group of ADP as well as the AlF_4_^−^ unit (Figure 6a) as concluded from the large ^31^P hyperfine coupling constant to the Pβ (a significantly smaller one is found for the Pα atom) and ^19^F and ^27^Al resonances observed in EDNMR, respectively. The structures of the only SF4-type helicases solved crystallographically, namely the *Bst*DnaB:GDP:AlF_4_^−^:DNA and *Aa*DnaB:ADP complexes (*Acquifex aeolicus*, PDB 4NMN^63^), support the finding of a Mn^2+^ coordination to the β-phosphate group as also supported by the DFT calculations of the ^31^P hyperfine tensors revealing the same trends as observed experimentally (Table S1, Figure S1). A similar experimental observation by EPR has been made for DbpA RNA helicase in complex with ADP^89^.

**Figure 6:**
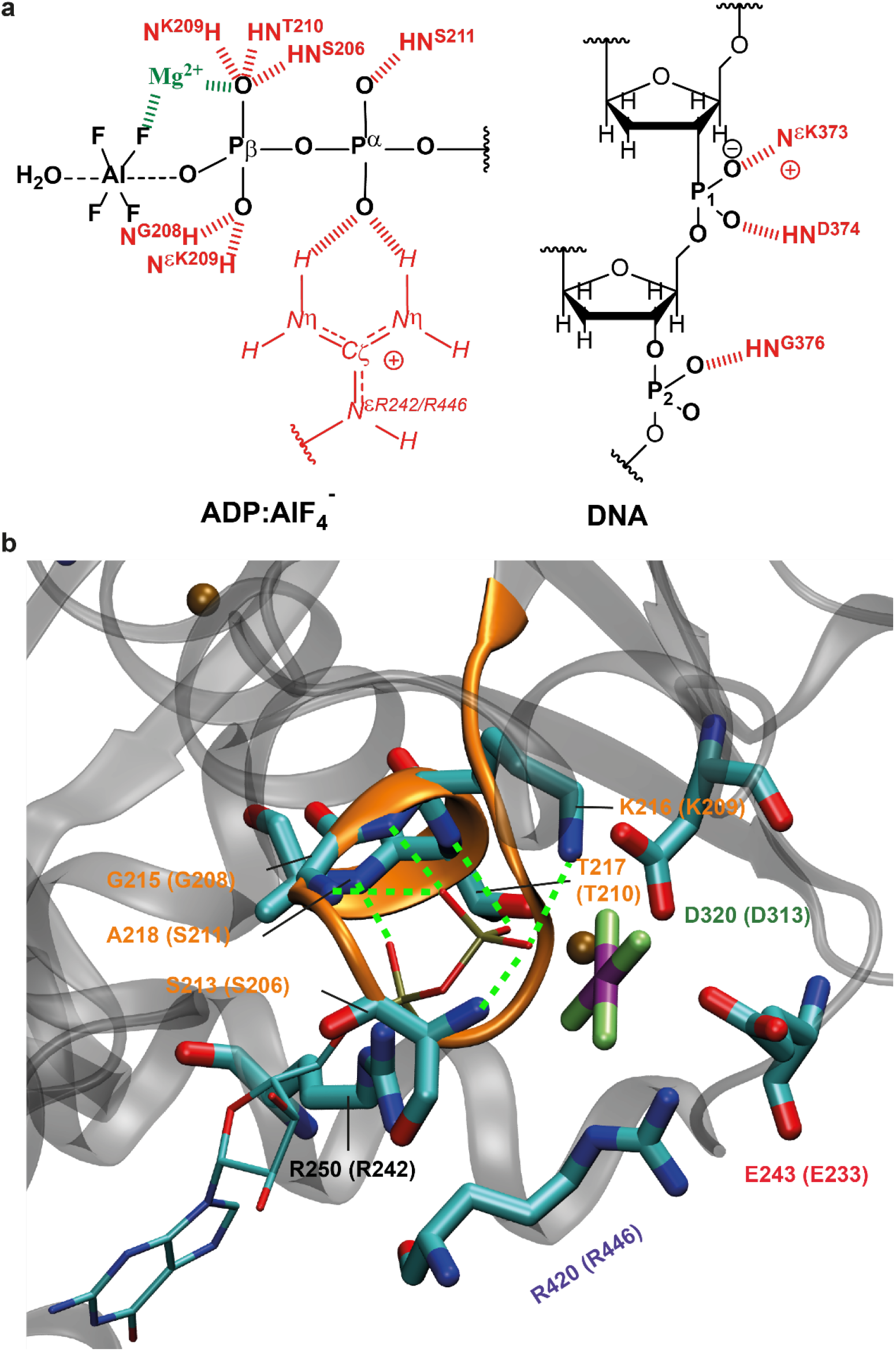
Comprehensive model for molecular recognition events involved in ADP and DNA binding to DnaB as obtained from the herein presented EPR/NMR results. **a** Sketch of hydrogen-bond formation and spatial proximities as revealed by the hPH and chemical-shift temperature-dependence experiments for ADP and DNA coordination to *Hp*DnaB. The Mg^2+^ cofactor has been placed in agreement with the results from the EDNMR spectra. **b** Zoom into the nucleotide binding domain for *Bst*DnaB:GDP:AlF_4_^−^:DNA (PDB accession code 4ESV). Residues given in brackets correspond to those in *Hp*DnaB. Green lines represent hydrogen bonds or spatial proximities as identified from the hPH correlation experiments.

Similar to other P-loop fold NTPases, in the *Hp*DnaB transition state trapped by solid-state NMR, residues S206, G208, K209, T210 and S211 of the Walker A motif were identified in coordinating the ADP phosphate groups by their backbone amino groups and by the side-chain of K209 yielding a dense hydrogen bond network (see Figure 6a for a schematic representation). DnaB helicases are characterized by a unique ARP[G/S]xGK[T/S] sequence of the Walker A motif with an Ala residue instead of Gly in the first position^16^. Homologous residues were found to coordinate the phosphate chain in the crystal structure of DnaB from *Bacillus stearothermophilus* (currently *Geobacillus stearothermophilus*) crystallized with Ca^2+^:GDP:AlF_4_^−^ and DNA (PDB accession code 4ESV^62^, see Figure 6b and Table S2 for the averaged distances to the oxygen atoms of the phosphate groups). An important difference is the only partial occupation of NBDs in *Bst*DnaB with the transition-state analogue, whereas for *Hp*DnaB all binding sites are occupied^66^ and highly symmetric as revealed by the absence of evident peak splitting in the hPH spectra.

The hPH spectrum reveals two key contacts in DNA recognition by DnaB, namely the coordination of D374 and G376 to the two structurally distinct DNA phosphate groups P1 and P2. The de-shielded proton resonances in combination with the almost vanishing temperature coefficient found for D374 point to an engagement of these two protons in hydrogen bonding (Figures 2 and 3). In previous studies, we have also identified the sidechain of K373 in forming a salt-bridge to P2^42^ which is also supported by the hPH spectrum showing a correlation of the resonance of the DNA phosphate group P2 to the K373 side-chain (Figure 2c). These contacts are identical to those found in the crystal structure of the *Bst*DnaB:DNA complex (backbone amide proton of E382 and G384 and the side-chain of R381)^62^ thus revealing similarities in DNA recognition for these two SF4-type helicases. The protein proton resonances contacting the DNA are not broadened or even split into several peaks indicating that all six DnaB subunits engage the DNA in a highly similar way, pointing to a closed hexamer rather than an extended open structure as observed for *Bst*DnaB (see Figure S9)^62^. This again agrees with our observation of a full saturation of all six NBDs with Mg^2+^:ADP:AlF_4_^−^ therefore still indicating structural differences in the position of DnaB monomers in the *Hp*DnaB and *Bst*DnaB helicase complexes with transition state analogues and DNA.

An important feature revealed in our NMR analysis is the free rotational diffusion of the AlF_4_^−^ unit mimicking the departing phosphate group during ATP hydrolysis. The averaging of the ^27^Al quadrupolar coupling constant in combination with the single ^19^F resonance observed indicate that the AlF_4_^−^ unit (Figure 4) is not coordinated tightly by the protein anymore, in contrast to the ADP for which we have observed a dense network of hydrogen bonds (Figures 2 and 3). Catalytic sites of oligomeric ATPases operate one after another so that catalysis in one subunit is thermodynamically promoted by substrate binding to the other subunit^90,91^. Hence, only one site stays at any moment in the conformation catalytically active for ATP hydrolysis. On the other hand, in contrast to the *Bst*DnaB structure, in the *Hp*DnaB complex studied herein ADP:AlF_4_^−^ moieties are present in all six catalytic pockets ^66^. The here reported free rotational diffusion of all six AlF_4_^−^ moieties within the tight hexamer of *Hp*DnaB (Figure 4) could be explained in the following way: The first binding of the ADP:AlF_4_^−^ moiety to a *Hp*DnaB subunit (subunit 1) brings it into its catalytically active, DNA-bound configuration with the Arg (R446) and Lys (K444) fingers of the adjoining subunit 2 interacting with the ADP:AlF_4_^−^:H_2_O_cat_ complex in the “catalytic” position. This suggestion is supported by our earlier observation that ADP:AlF_4_^−^ binding alone induces protein conformational changes and preconfigures the protein for DNA binding^66^. This re-arrangement might be driven by the exergonic AlF_4_^−^ binding in the catalytic site, as discussed elsewhere^19^. Binding of the ADP:AlF_4_^−^:H_2_O_cat_ to the subunit 2 transforms it in a similar way and, simultaneously, provides free energy for pulling the γ-phosphate-mimicking AlF_4_^−^ out of its catalytic position in the subunit 1 by Arg and Lys fingers of subunit 2. After this sequence of events repeats six times, all six protein subunits are in the same catalytic configuration being tightly fixed on the DNA strand (as revealed by the identified hydrogen bonds formed by D374 and G376 to the DNA phosphate groups, Figure 2c) whereas their six AlF_4_^−^ moieties are in positions similar to those taken by AlF_4_^−^ moieties in two of six catalytic sites of *Bs*tDnaB, namely those on the subunit interfaces B/C and F/A (Figure 5c, S8). In this state, AlF_4_^−^ moieties are detached both from the ADP moiety and the Arg and Lys fingers. Hence, we suggest that the mobile AlF_4_^−^ moiety in *Hp*DnaB mimics the cleaved phosphate group during the late transition state of ATP hydrolysis by *Hp*DnaB; at this stage, the phosphate unit is nearly released from the NBD. In this situation, when ATP hydrolysis has already proceeded, the protein could use the chemical energy released for DNA translocation.

Our results demonstrate that magnetic resonance is highly suitable to obtain structural and dynamic insights into the transition state of ATP hydrolysis of a bacterial DnaB helicase trapped by aluminum fluoride allowing a more profound understanding of the functioning of such complex motor proteins. EPR reveals the coordination of the metal ion co-factor to the β-phosphate group of ADP as well as to the AlF_4_^−^ unit, whereas proton-detected hPH solid-state NMR experiments combined with temperature-dependences of proton chemical-shift values allow for identifying hydrogen bonds which are crucial for the molecular recognition process of ADP and DNA binding to the DnaB helicase. NMR is one of the most sensitive techniques in proving hydrogen bonding with the additional advantage of shedding light onto dynamic processes, herein the free rotational diffusion of the AlF_4_^−^ unit mimicking the phosphate group transferred during ATP hydrolysis.

## Methods

### Sample preparation

#### Protein expression and purification

Natural abundance and ^13^C-^15^N labelled *Hp*DnaB was prepared in buffer A (2.5 mM sodium phosphate, pH 7.5, 130 mM NaCl) as described in reference ^51^. In short, DnaB was recombinantly expressed in presence of ^13^C-glucose (2 g/L) and ^15^N-ammonium chloride (2 g/L) as sole sources of carbon-13 and nitrogen-15. In case of the deuterated protein, the protein was expressed in D_2_O in presence of deuterated ^13^C-glucose. The back-exchange was achieved by purifying the protein in a protonated buffer (2.5 mM sodium phosphate, pH 7.5, 130 mM NaCl).

#### NMR sample preparation

0.3 mM *Hp*DnaB in buffer A was mixed with 5 mM MgCl_2_ ∙ 6H_2_O and consecutively 6 mM of an NH_4_AlF_4_ solution (prepared by incubating 1 M AlCl_3_ solution with a 5-fold excess of 1M NH_4_F solution (compared to AlCl_3_) for 5 min) and 5 mM ADP and incubated for 2 h at 4°C. 1 mM of (dT)_20_ (purchased from Microsynth) was added to the complexes and reacted for 30 min at room temperature. The protein solution was sedimented^51,52,92^ into the MAS-NMR rotor (16 h at 4°C at 210’000 *g*) using home-built tools^93^. In case of the DnaB:ADP:AlF_4_^−^ complex the DNA addition step was omitted.

#### EPR sample preparation

For EPR experiments, natural abundance DnaB was concentrated to 48 mg/ml (850 μM) using a Vivaspin 500 centrifugal filter with a cut-off of 30 kDa. The concentrated protein was incubated in presence of 6 mM ADP, 170 μM Mn^2+^ and 7 mM NH_4_AlF_4_ for 2 h at 4°C. After 2 h, glycerol was added to a concentration of 20 %. The final concentrations were: DnaB 690 μM, ADP 5 mM, Mn^2+^ 138 μM, and NH_4_AlF_4_ 6 mM.

### Solid-state NMR experiments

Solid-state NMR spectra were acquired at 11.7, 14.1 and 20.0 T static magnetic-field strengths using an in-house modified Bruker 3.2 mm (^19^F and ^27^Al NMR) probe and a 0.7 mm (^1^H NMR) triple-resonance (^1^H/^31^P/^13^C) probe. The MAS frequencies were set to 17 and 100/105 kHz, respectively. The 2D spectra were processed with the software TOPSPIN (version 3.5, Bruker Biospin) with a shifted (2.0 or 3.0) squared cosine apodization function and automated baseline correction in the indirect and direct dimensions. For ^1^H-detected experiments, the sample temperature was set to 293 K^93^ and varied in the range of 294-302 K for the temperature dependence studies. A fast adjustment of the temperature in the bore of the magnet (typically causing *B*_0_ instabilities) was achieved by a bore heating system implemented by the instrument manufacturer. This is crucial for detecting the rather small temperature dependences of proton chemical-shift values (on the order of several ppb/K)^58^. For ^19^F (recorded at 14.1 T) and ^27^Al (recorded at 11.7 T) MAS-NMR experiments, the sample temperature was adjusted to 278 K. ^1^H and ^31^P-detected spectra were analysed with the software CcpNmr^94–96^ and referenced to 4,4-dimethyl-4-silapentane-1-sulfonic acid (DSS). ^19^F and ^27^Al spectra were referenced to internal standards. For more detail see Table S3 in the Supplementary Materials Section.

### EPR experiments

All experiments were conducted on a Bruker Elexsys E680 EPR spectrometer (Bruker Biospin) operating at W-band frequencies (approximately 94.2 GHz). ENDOR measurements used a 250 W radiofrequency (rf) amplifier. The temperature was generally set to 10 K.

Electron-electron double resonance (ELDOR)-detected NMR spectra were acquired with a shot repetition time of 1 ms and the echo-detected hole-burning sequence t_HTA_ – T – t_p_ – τ – 2t_p_ – τ – echo, with t_HTA_= 50 μs, T = 10 μs, t_p_ = 100 ns, τ = 1400 ns and an integration window of 1400 ns. The frequency of the high-turning angle (HTA) pulse was incremented in steps of 0.1 MHz over the measured range. A +/− phase cycle on the first π/2 pulse of the echo was used to eliminate unwanted coherence transfer pathways. The power of the HTA pulse, generated by the ELDOR channel of the spectrometer, was optimized such that the observed lines were as intense as possible without being broadened by saturation effects. The nutation frequency ν_1_ at the centre of the resonator was about 6 MHz. The settings were held constant between protein samples and the corresponding control samples. Yet it is important to note that exact reproducibility of peak intensities between runs may be difficult with the resonator used because the resonator profile strongly affects line intensities in EDNMR, and hence a careful experimental setup is required.

Davies ENDOR spectra were acquired with a shot repetition time of 5 ms and with the sequence t_inv_ – T – t_p_ – τ – 2t_p_ – τ – echo, where during the time T, an rf-pulse was applied. The inversion pulse length was set to 200 ns, and the rf-pulse length to 50 μs. The echo was integrated symmetrically around the echo maximum over a time of 400 ns. Due to enormous time overhead on this particular spectrometer, we did not use stochastic acquisition mode and used 10 shots per point.

Mims ENDOR spectra were acquired with a shot repetition time of 2.5 ms and with the sequence t_p_ – τ – t_p_ – T – t_p_ – τ – echo, where during the time T, an rf-pulse was applied. The interpulse delay τ was set to 1200 ns, corresponding to the phase memory time T_m_, where detection of small hyperfine couplings is most sensitive^97^, and the rf-pulse length to 25 μs.

Raw EDNMR data were background corrected with a Lorentzian line that was fitted to the central hole, and normalized to the signal intensity far off-resonance, i.e. the peak intensity corresponds to the relative hole depth.

### DFT calculations of hyperfine tensors

DFT calculations were performed on small clusters mimicking the coordination sphere of the metal ion co-factor extracted from the PDB structures (*Bst*DnaB: accession code 4ESV and *Aa*DnaB: accession code 4NMN, see Figure S1). Hydrogen atoms were added to saturate terminating groups and their positions were optimized on a TPSS^98^/def2-SVP^99^ level using TURBOMOLE (version 6.0)^100,101^. In all TURBOMOLE SCF calculations, an energy convergence criterion of 10^−7^ E_h_ and in all geometry optimizations an energy convergence criterion of 5×10^−7^ E_h_ was chosen. The integration grid was set to m4 and the RI approximation was used. Hyperfine coupling tensors were calculated in the ADF suite (version 2013)^102^ on a B3LYP^103,104^/TZ2P^105^ level of theory. The INTEGRATION keyword was set to 6.0 and in the SCF calculation an energy convergence criterion of 10^−6^ E_h_ was used.

### Data Availability

Data supporting the findings of this manuscript are available from the corresponding authors upon reasonable request. The following PDB structures were used in this study: 4ZC0, 4NMN, 4ESV, 3CMW and 3PUW. All experimental NMR parameters are provided as a Source Data file.

## Acknowledgements

This work was supported by the ETH Career SEED-69 16-1 (T.W.) and the ETH Research Grant ETH-43 17-2 (T.W.), the Deutsche Forschungsgemeinschaft (DFG, German Research Foundation, project number 455240421 and Heisenberg fellowship, T.W.), an ERC Advanced Grant (B.H.M., grant number 741863, Faster) and by the Swiss National Science Foundation (B.H.M., grant number 200020_159707 and 200020-188711), the German Academic Exchange Service (DAAD, M.I.K.) and the EvoCell Program of the Osnabrueck University (A.M.Y). T.W. acknowledges discussions with Prof. Matthias Ernst and Dr. Denis Lacabanne.

## Author contributions

AAM, LAV and TW performed the NMR experiments, NW and DK the EPR experiments. RC prepared the samples. AD modified the NMR probes for ^27^Al and ^19^F experiments. TW performed the DFT calculations. MIK and AYM performed the structural modellings. AAM, LAV, NW, DK, MEW, JZ, HE, GJ, AYM, AB, BHM and TW analyzed the data. DK, AYM, BHM and TW designed and supervised the research. All authors contributed to the writing of the manuscript.

