## Supplementary Material for "Spectroscopic glimpses of the transition state of ATP hydrolysis trapped in a bacterial DnaB helicase"

### Supplementary Figures and Tables

**Table S1**: Experimental and calculated (in brackets) $^{31}$P hyperfine coupling tensor values. The models used for the DFT calculations are shown in Figure S1. δ denotes the anisotropy and $\eta_A$ the asymmetry of the hyperfine coupling tensor. [a] calculated values based on the *Bst*DnaB model [b] calculated values based on the *Aa*DnaB model.

| | $A_{iso}$/ MHz | $A_{xx}$/ MHz | $A_{yy}$/ MHz | $A_{zz}$/ MHz | δ/ MHz | $\|\eta_A\|$ |
|---|---|---|---|---|---|---|
| $^{31}$Pα | 0.3 (0.05[a]/0.01[b]) | 0.23 (-0.24[a]/ -0.32[b]) | 0.23 (-0.23[a]/ -0.28[b]) | 0.37 (0.61[a]/0.64[b]) | 0.14 (0.85[a]/ 0.66[b]) | 0.00 (0.02[a]/ 0.06[b]) |
| $^{31}$Pβ | 4.7 (3.35[a]/12.8[b]) | 3.8 (2.36[a]/11.3[b]) | 3.8 (2.63[a]/11.5[b]) | 6.4 (5.07[a]/15.6[b]) | 2.6 (2.58[a]/ 4.20[b]) | 0.00 (0.16[a]/ 0.07[b]) |

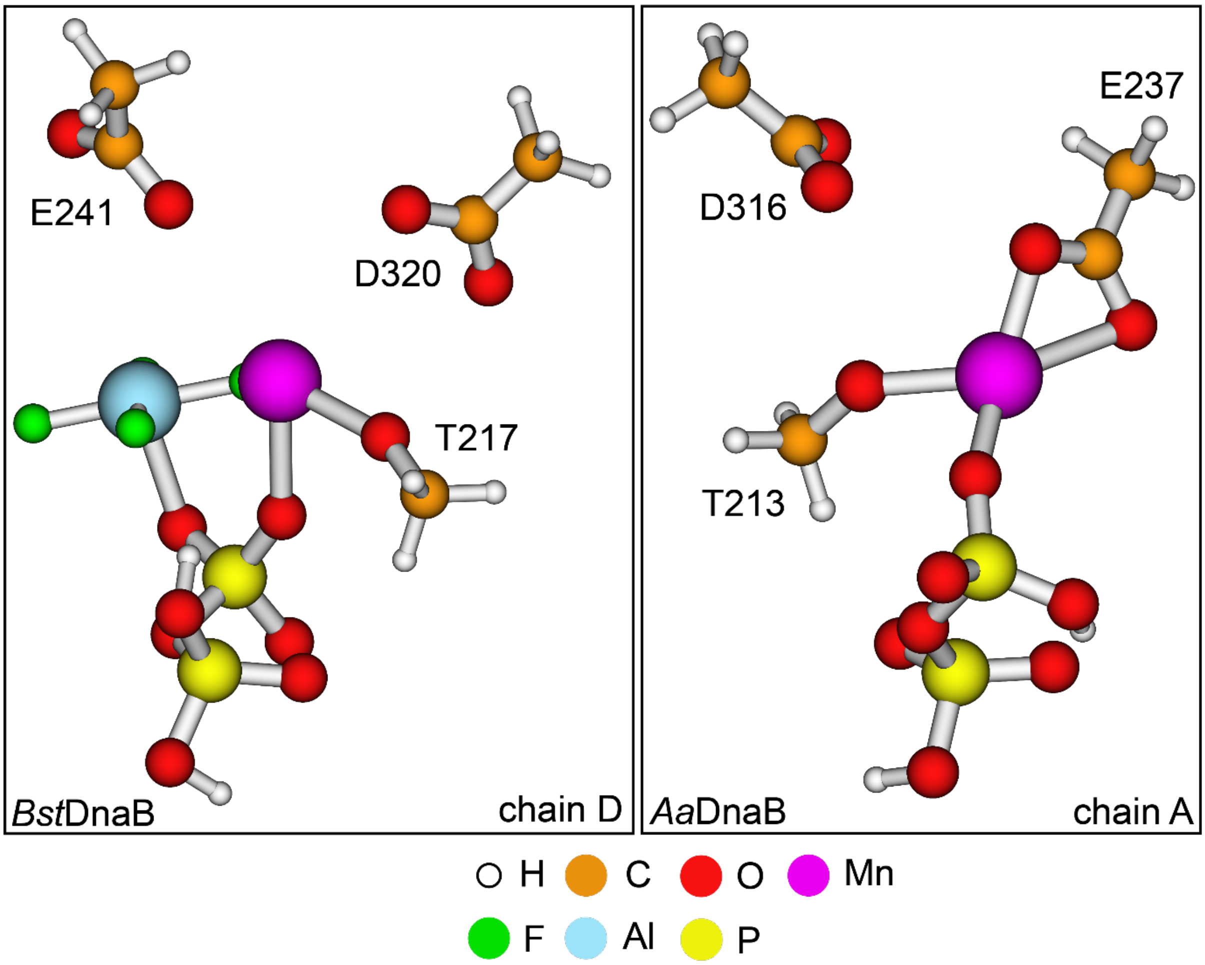


**Figure S1**: Models extracted from the PDB files 4ESV (*Bst*DnaB) and 4NMN (*Aa*DnaB) used in the DFT calculations of the hyperfine coupling tensors.

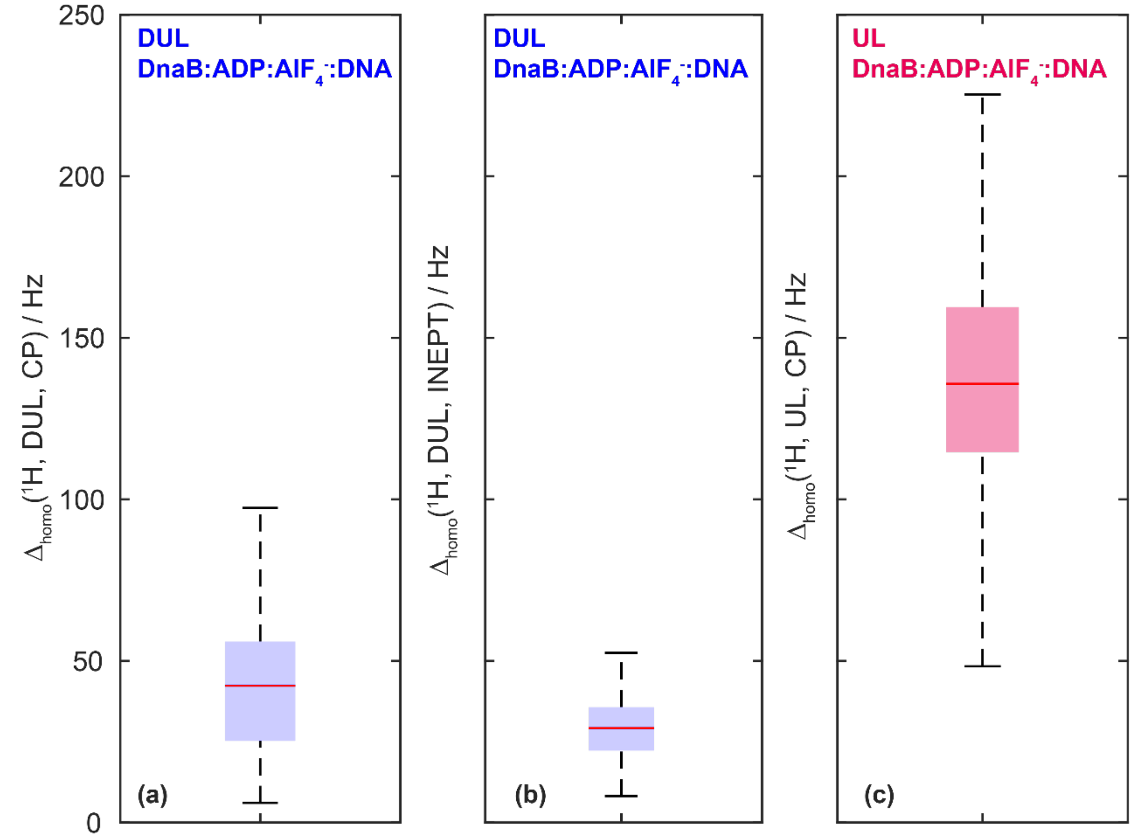


**Figure S2:** *The proton-linewidths of the deuterated and 100 % back-exchanged sample decrease compared to a fully-protonated sample.* Boxplot representation of homogeneous proton linewidth statistics in DnaB:ADP:$AlF_4^-$:DNA determined from site-specific $T_2$'($^1$H) measurements on isolated resonances in a 2D hNH spectrum of a deuterated and 100 % back-exchanged sample ((**a**) using CP and (**b**) using refocused INEPT as a polarization transfer mechanism) and for a fully protonated sample (**c**). Latter have already been reported in reference[1]. The homogeneous proton line-widths roughly decrease by a factor of three in the deuterated compared to the fully-protonated sample.

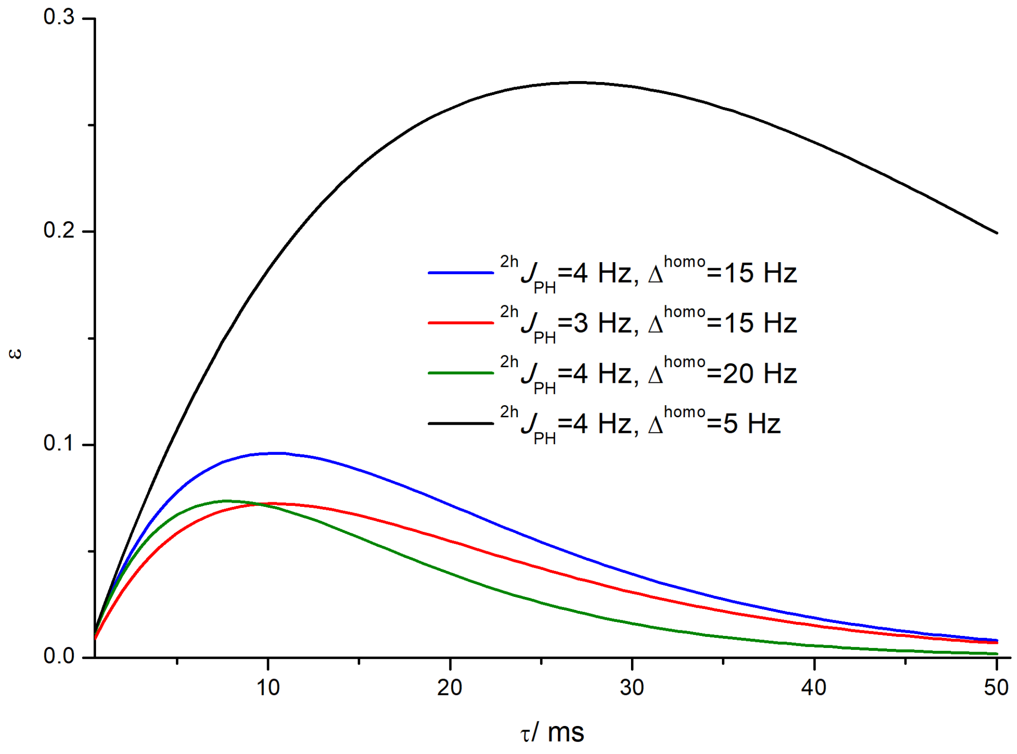


**Figure S3**: INEPT enhancement factor ε as a function of the mixing time assuming different (relatively small) $\Delta^{homo}$ values and $^{2h}J_{PH}$ J-coupling constants. Infinitely long $^{31}P$ transverse relaxation has been assumed for all simulations. The simulations have been performed based on the well-known formulas for an HX spin system[2-4]. The $^{2h}J_{PH}$ couplings are so weak that even for the smallest homogeneous linewidth of 5 Hz, the theoretical transfer efficiency stays below 30%, while it is already less than 10% in all other cases.

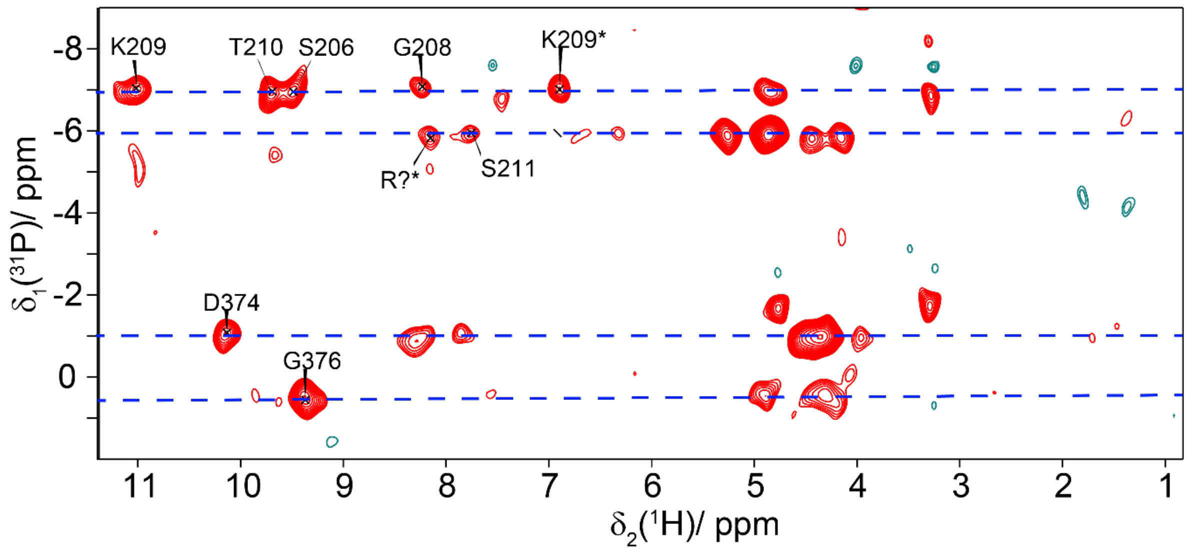


**Figure S4**: hPH correlation spectrum recorded at 20.0 T with a MAS frequency of 105 kHz and using a forward and backward CP-contact time of 1.5 ms. The dashed blue lines indicate the chemical-shift positions of the four $^{31}P$ resonances from Figure 2a. * stands for correlations to side-chain protons.

| Residue | Average distance/ Å |
|---|---|
| N-O3B S213 (S206) | 3.0 |
| N-O2B G215 (G208) | 3.1 |
| N-O2B K216 (K209) | 2.4 |
| NZ-O3B K216 (K209) | 2.7 |
| N-O1B T217 (T210) | 3.3 |
| N-O1A/O2A A218 (S211) | 3.1 |

**Table S2**: Distances between the oxygen phosphate groups of GDP and protein nitrogen atoms in the *Bst*DnaB crystal structure (PDB:4ESV). The residues shown in brackets correspond to the ones of *Hp*DnaB. The average distance for the five bound GDP molecules is given. Note, that for A218 the closest distance to one of the two oxygen atoms O1A and O2A has been chosen.

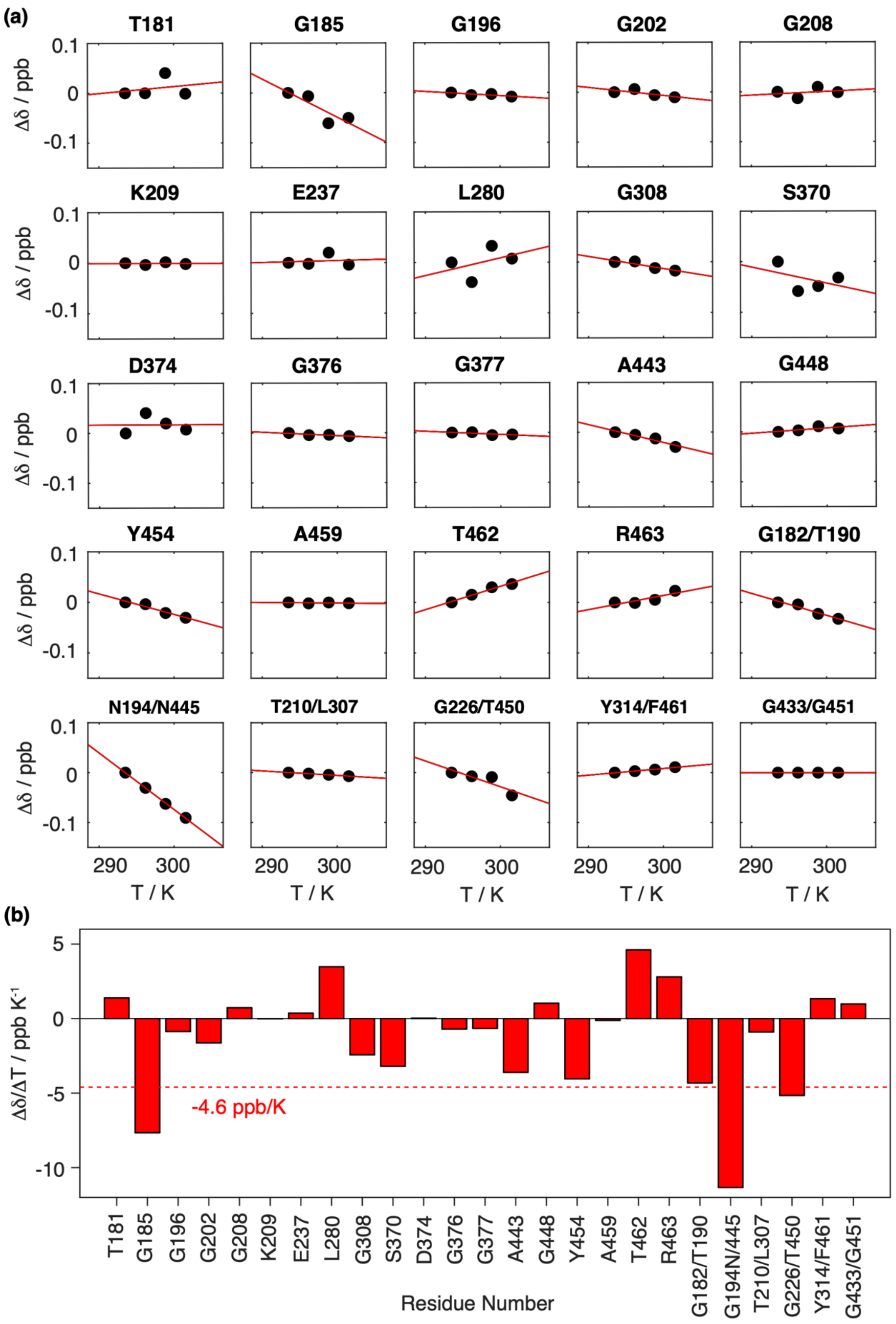


**Figure S5**: *Proton chemical-shift temperature coefficients.* **(a)** Residue-specific temperature-dependent proton chemical-shift values (black circles) between 294-302 K and linear fit (red) for the extraction of correspondent temperature coefficients. The chemical shifts are referenced to the corresponding value at 294 K. **(b)** Site-specific proton chemical-shift temperature coefficients determined from the slope of the linear regression of the data points shown in **(a)**.

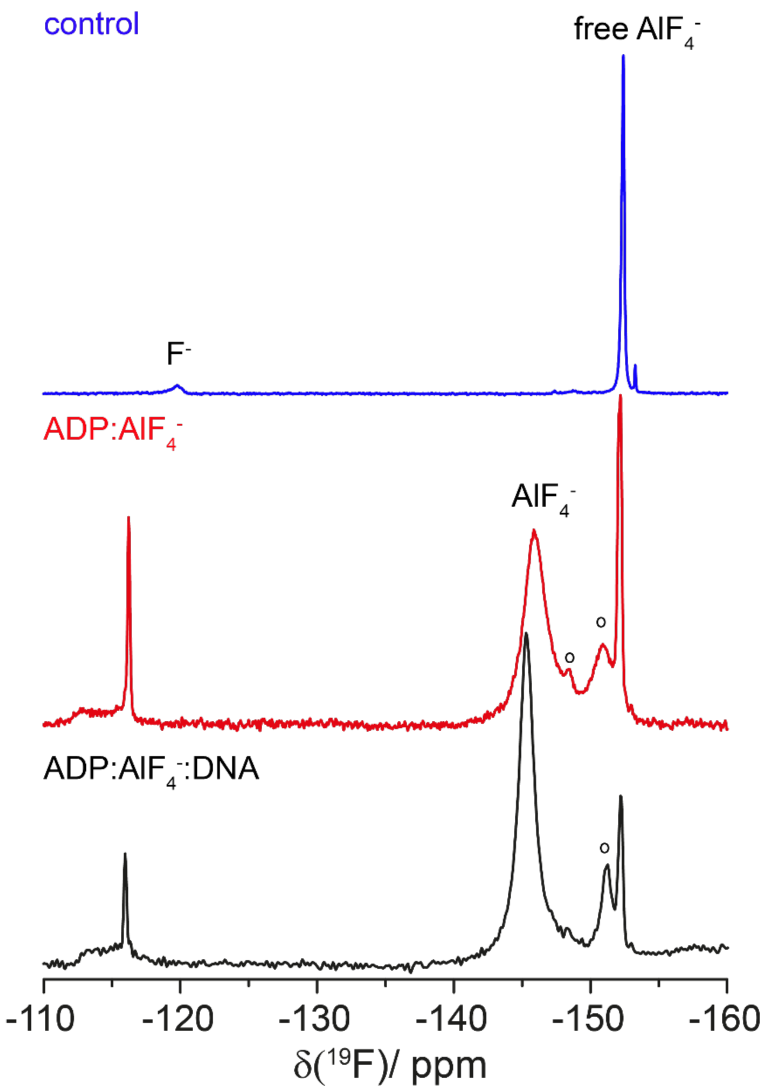


**Figure S6:** *$^{19}$F MAS spectra indicate the fast rotation of the AlF$_x$ unit.* $^{19}$F MAS spectra recorded at 14.1 T with an MAS frequency of 17.0 kHz and with the EASY background suppression scheme[5]. Spectra were acquired on DnaB:ADP:AlF$_4^-$ in the presence and absence of DNA, as well as on the buffer solution without protein (control experiment). o indicates precipitated AlF$_x$(OH)$_{6-x}$ species.

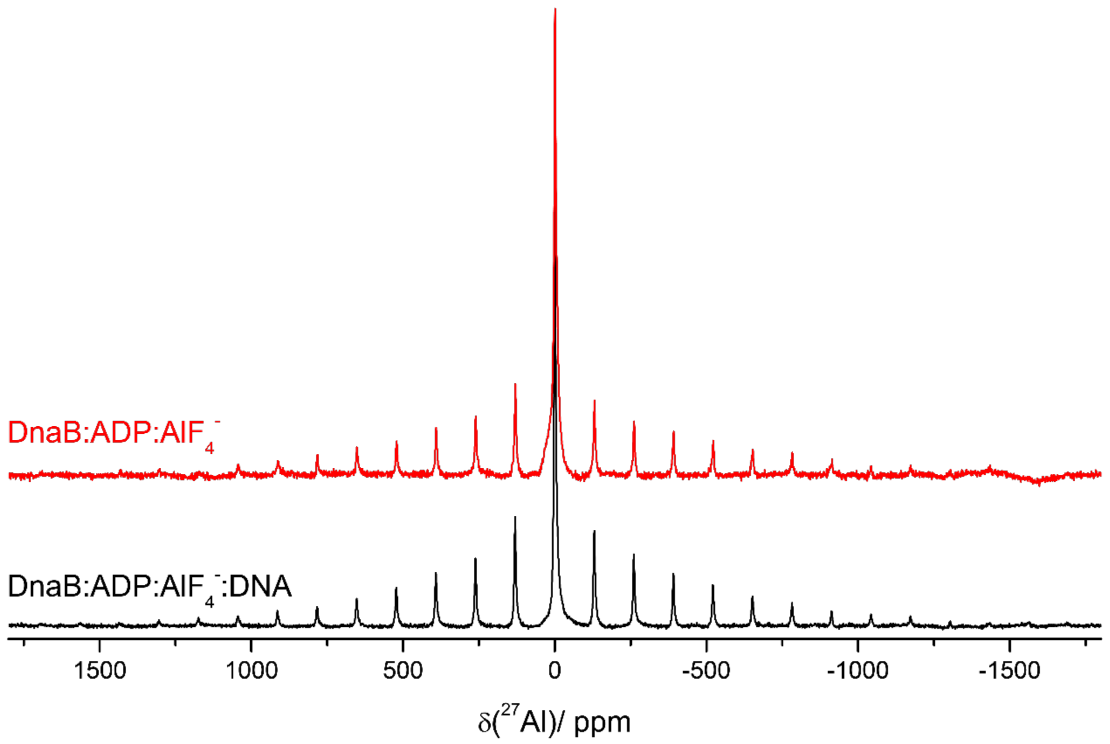


**Figure S7**: *The AlF$_4^-$ unit is mobile in absence and presence of DNA.* $^{27}$Al MAS spectra of DnaB:ADP:AlF$_4^-$ and DnaB:ADP:AlF$_4^-$:DNA recorded at 11.7 T and 17 kHz MAS.

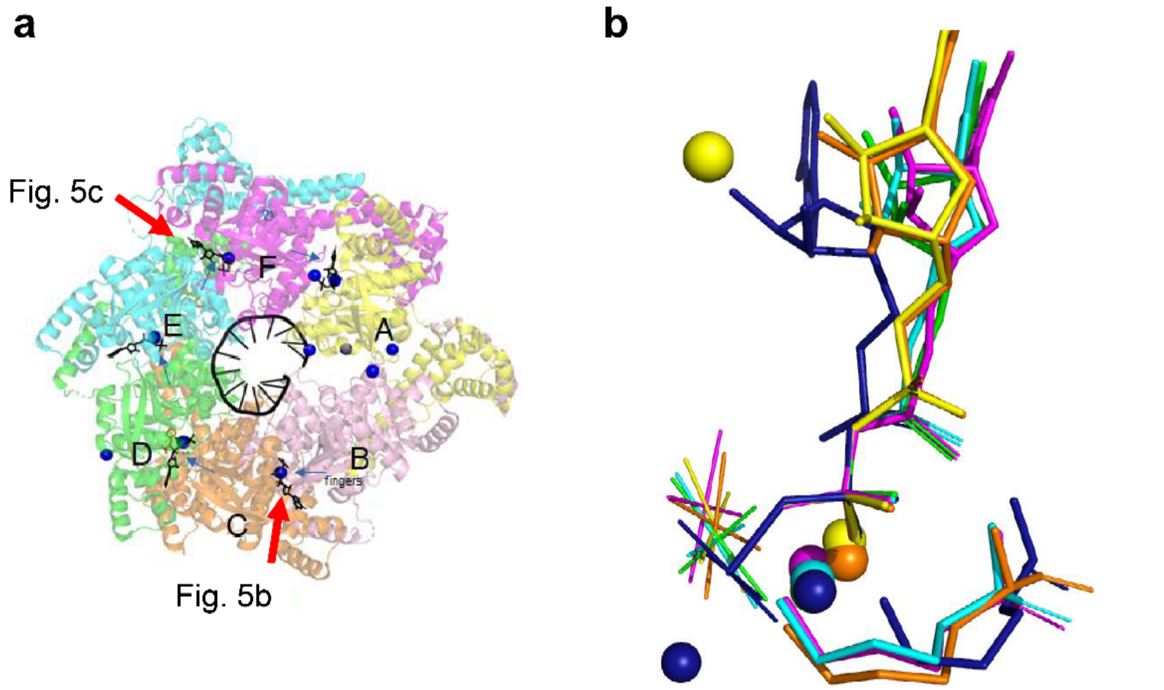


**Figure S8**: Relative orientation of the $AlF_4^-$ moieties in all five occupied binding sites of the DnaB structure from *Geobacillus stearothermophilus* (*Bst*DnaB, see also Figure 5, PDB accession code 4ESV and reference [6]). The colors of $AlF_4^-$ moieties (**b**) correspond to the colors of subunits to which they are bound (**a**). For comparison, the dark-blue colored structure of the ADP:$AlF_4^-$ : $H_2O_{cat}$ complex from the ABC ATPase of the maltose transporter MalK (see PDB accession code 3PUW and reference [7]) is shown. NDPs were superimposed by atoms $O^{3A}$, $P^B$ and $O^{3B}$ in Pymol[8].

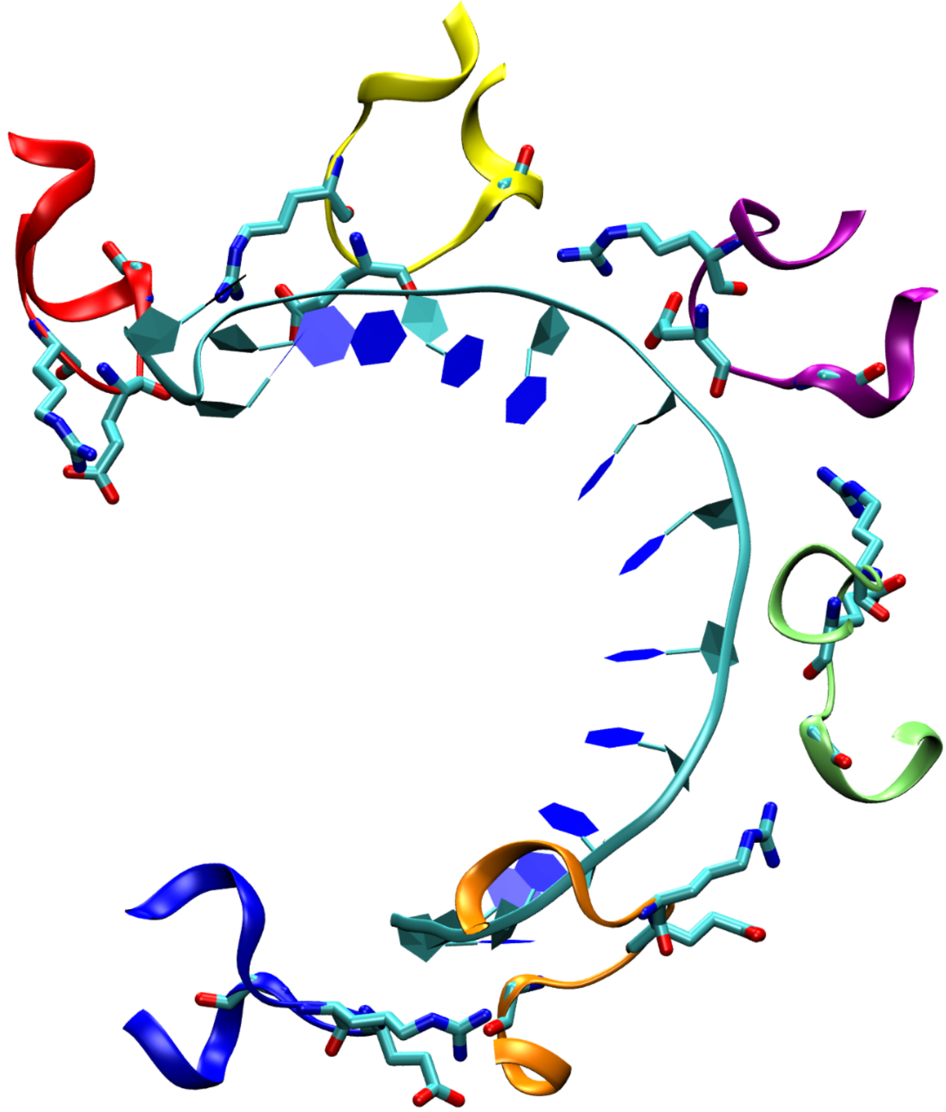

**Figure S9**: *DNA binding in BstDnaB.* DNA binding in the *Bst*DnaB:DNA complex (PDB 4ESV). The DNA binding loops (residues 378-387) are shown in different colors. Residues R381, E382 and G384 are shown in licorice style. The DNA is shown in ribbon representation.

**Table S3**: Experimental solid-state NMR parameters.

| **Experiment** | **hPH (DUL DnaB:ADP:AlF$_4^-$:DNA)** | **hPH (DUL DnaB:ADP:AlF$_4^-$:DNA)** | **hNH** (DUL DnaB:ADP:AlF$_4^-$:DNA)** |
|---|---|---|---|
| $\nu_r$/ kHz | 105 | 105 | 100 |
| $B_0$/ T | 20.0 | 20.0 | 20.0 |
| **Transfer I** | HP-CP (DQ) | HP-CP (DQ) | HN-CP(ZQ) |
| $\nu_1$($^1$H)/ kHz | 65* | 65* | 146 |
| $\nu_1$($^{15}$N/$^{31}$P)/ kHz | 34.4* | 35.6* | 31 |
| CP contact time/ ms | 3.5 | 1.5 | 0.8 |
| Shape | Tangent $^1$H | Tangent $^1$H | Tangent $^1$H |
| **Transfer II** | PH-CP (DQ) | PH-CP (DQ) | NH-CP(ZQ) |
| $\nu_1$($^1$H)/ kHz | 65* | 65* | 146 |
| $\nu_1$($^{31}$P/$^{15}$N)/ kHz | 34.4* | 35.6* | 31 |
| CP contact time/ ms | 3.5 | 1.5 | 1.2 |
| Shape | Tangent $^1$H | Tangent $^1$H | Tangent $^1$H |
| $^1$H carrier/ ppm | 4.8 | 4.8 | 4.8 |
| $^{31}$P/$^{15}$N carrier/ ppm | -0.6 | -0.6 | 117.5 |
| $t_1$ increments | 128 | 128 | 200 |
| Sweep width ($t_1$)/ ppm | 40 | 40 | 70 |
| Acquisition time ($t_1$)/ ms | 4.6 | 4.6 | 16.6 |
| $t_2$ increments | 1024 | 768 | 2048 |
| Sweep width ($t_2$)/ ppm | 46.7 | 46.7 | 46.7 |
| Acquisition time ($t_2$)/ ms | 12.9 | 9.7 | 25.8 |
| water Suppression | MISSISIPPI | MISSISIPPI | MISSISIPPI |
| $\nu_1$($^1$H) / kHz | 30 | 30 | 20 |
| Time water suppression / ms | 120 | 120 | 120 |
| $^{15}$N/$^{31}$P WALTZ64 decoupling power/ kHz | 5 | 5 | 5 |
| Interscan delay/ s | 1 | 1 | 1.2 |
| Number of scans | 3232 | 3968 | 224 |
| Measurement time/ h | 115 | 141 | 17 |

* The CP conditions were optimized on a solid sample of ortho-phospho-L-serine.

** The experiments were performed at sample temperatures from 294 to 302 K within one measurement session and the same experimental parameters.

**Table S3**: Experimental solid-state NMR parameters (continued).

| **Experiment** | **$^{27}Al$ (UL DnaB:ADP:$AlF_4^-$ :DNA)** | **$^{19}F$ (UL DnaB:ADP:A lF$_4^-$:DNA)** | **$^{19}F$ (UL DnaB:ADP:$AlF_4^-$)** | **$^{19}F$ (control)*** ** |
|---|---|---|---|---|
| $\nu_r$/ kHz | 17.0 | 17.0 | 17.0 | 17.0 |
| $B_0$/ T | 11.7 | 14.1 | 14.1 | 14.1 |
| **Type** | $^{27}Al$ direct | $^{19}F$ EASY[5] | $^{19}F$ EASY[5] | $^{19}F$ EASY[5] |
| $\nu_1(^{19}F)$ / kHz | - | 50** | 50** | 50** |
| $\nu_1(^{27}Al)$ / kHz | 33* | - | - | - |
| $t_1$ increments | 4096 | 8192 | 8192 | 8192 |
| Sweep width ($t_1$)/ ppm | 4795 | 295 | 295 | 295 |
| Acquisition time ($t_1$)/ ms | 3.3 | 24.6 | 24.6 | 24.6 |
| Interscan delay/ s | 1.5 | 5.0 | 5.0 | 5.0 |
| Number of scans | 40960 | 1536 | 1536 | 1024 |
| Measurement time/ h | 17 | 2.1 | 2.1 | 1.4 |

* : Determined from the nutation frequency of the measured sample. A 30° excitation pulse was used. The $^{27}Al$ spectra were referenced to $Al(NO_3)_3$ using solid $(NH_4)Al(SO_4)_2$ resonating at -0.4 ppm.

**: Determined from the nutation frequency of the measured sample. The $^{19}F$ spectra were referenced relative to $CFCl_3$ using solid $NaBF_4$ resonating at -159.2 ppm.

***: Control solution using the same buffer than for the protein and the same ADP, $NH_4AlF_4$ and $Mg^{2+}$ concentrations.
